# kamino: fast proteome-wide variant calling for amino acid phylogenomics

**DOI:** 10.64898/2026.05.21.726148

**Authors:** Romain Derelle, John A. Lees, Leonid Chindelevitch

## Abstract

Amino acid-based phylogenetics usually relies on first clustering and aligning orthologous proteins. This approach is powerful but computationally demanding. Here, we present kamino, a reference- and alignment-free method that rapidly builds amino acid phylogenomic alignments directly from proteomes. As with similar algorithms, homologous regions are identified through shared sequences flanking variable regions. The method uses local changes in recoded k-mer occupancy to efficiently identify variable positions within these homologous regions and extract the corresponding pseudo-aligned sequences. It generates phylogenetically informative alignments across diverse prokaryotic and eukaryotic datasets. Phylogenetic analyses show that it accurately recovers Mycobacterium tuberculosis lineages, most curated GTDB taxa, and relationships consistent with published Drosophila and mammalian phylogenies, while producing signals broadly similar to BUSCO-based approaches. Runtimes are comparable to genome-based alignment-free methods and several orders of magnitude faster than classical marker-based pipelines, with moderate memory requirements. The method performs well across a broad range of divergence levels, from within-species comparisons to family-level prokaryotic and phylum-level eukaryotic datasets. kamino therefore provides a fast and simple route from proteomes to phylogenomic alignments across a broad range of evolutionary scales. The program is implemented in Rust and freely available at https://github.com/rderelle/kamino.

## Introduction

Phylogenomic analyses across distant evolutionary relationships typically rely on multi-gene amino acid alignments (Rokas et al. 2003; Delsuc et al. 2005; Kapli et al. 2020). This involves clustering all proteins from the studied species into orthogroups, then filtering to retain reliable phylogenetic markers based on factors such as paralogy and missing data. This approach can generate large, tailored datasets, but requires computationally intensive all-vs-all protein clustering (Cosentino et al. 2024; Klemm et al. 2023; Emms et al. 2025) and careful curation of gene families to identify paralogy events, making it time-consuming and difficult to scale. Reduced sets of conserved phylogenetic markers, typically single-copy genes present across the dataset, offer a faster but lower-resolution alternative to full orthogroup inference (Simão et al. 2015; Tegenfeldt et al. 2025). These markers can be identified in each proteome using similarity searches such as HMMER (Eddy 2011) or DIAMOND (Buchfink et al. 2021), allowing scalable analyses across much broader taxon sampling (Segata et al. 2013; Na et al. 2018; Kim et al. 2023; Holmes and Kelly 2026). Alternatively, faster alignment-free approaches based on amino acid k-mer distributions have been proposed (Leimeister et al. 2019; Tessa Pierce-Ward et al. 2022). However, these methods only produce pairwise distances, making them less suitable for accurate phylogenetic inference across deep evolutionary divergences.

Some of the most important recent advances in fast alignment for phylogenetics have come from methods designed for within-species genomic analyses. Beyond approaches that estimate inter-genome distances directly (Ondov et al. 2016; Jain et al. 2018; von Wachsmann et al. 2025), several algorithms generate nucleotide alignments by identifying homologous positions without constructing a whole-genome alignment or requiring a reference genome. Two strategies have been used: the split k-mer method matches mutations by identifying conserved sequence flanking a variable middle base (Derelle et al. 2024; Hall and Nisbet 2023), whereas assembly graph approaches identify bubbles (i.e., alternative paths between shared graph nodes) that represent variable sites or clusters of nearby variable sites (Uricaru et al. 2015; Derelle et al. 2025; Iqbal et al. 2012). In both cases, conserved sequence context serves as a local reference for inferring homology, allowing genomes to be decomposed on the fly into sets of homologous positions from which variants can be called efficiently. These approaches also avoid explicit orthology/paralogy assignment by collapsing homologous sequences from the same isolate between shared anchors into a strict consensus. The key limitation is that these conserved flanking sequences must be shared across genomes; as genomic diversity increases, fewer positions can be confidently matched, confining such approaches largely to population- or species-level studies. High genomic diversity can also substantially increase memory requirements, as these methods retain all observed k-mers or graph elements in memory.

We adapted this principle of local homology inference to proteome data, generating amino acid alignments for phylogenetics across a broader range of evolutionary divergences. The resulting program, kamino, builds a phylogenomic alignment directly from a set of proteomes, without first identifying orthologous gene families or aligning complete proteins. Instead, kamino uses an alignment-free approach to group homologous variable sequences into variant groups, defined as sets of pseudo-aligned sequences flanked by the same pair of conserved sequences and supported across most proteomes (Derelle et al. 2025). It extracts these variant groups in two passes.

The first recodes amino acids into a six-state alphabet, by default SR6 (Susko and Roger 2007), which collapses the 20 amino acids into six characters by grouping chemically similar residues that commonly substitute for one another, then estimates the occupancy of the recoded k-mers across all proteomes. The second uses local changes in that occupancy to identify conserved sequences flanking short variable regions, then extracts them in the original 20-amino-acid alphabet. Rather than storing every observed k-mer explicitly, kamino uses compact probabilistic summaries of k-mer occupancy and only retains the extracted variant groups. This keeps runtime and memory usage relatively low. Below, we describe the algorithm and evaluate its performance across diverse proteome datasets. We show that kamino produced informative alignments with fast runtimes and modest memory use, recovered the monophyly of most curated bacterial taxa tested, and yielded phylogenetic signals broadly similar to marker-based approaches.

## Materials and methods

### Algorithm description

kamino takes as input one proteome FASTA file per sample. Files can be provided directly from a directory or through a two-column table containing sample names and proteome paths, and can be gzip-compressed. Each protein is analysed independently, and regions containing ambiguous amino acids are excluded from k-mer matching.

Variant groups are extracted in two passes over the proteomes (Figure 1). In the first pass, amino acid sequences are converted to the selected six-state alphabet and decomposed into overlapping k-mers (k=13 by default). kamino estimates the occupancy of each recoded k-mer, defined as the number of input proteomes in which it occurs. A Bloom filter for each proteome prevents repeated occurrences of the same k-mer from being counted more than once, while a global count-min sketch tracks the resulting k-mer occupancies without retaining all k-mers explicitly. In the second pass, kamino scans each recoded protein again for local decreases and increases in k-mer occupancy. A k-mer is considered shared when it occurs in at least ⌈f × N⌉ of the N input proteomes, where f is the minimum sample occupancy (f=0.85 by default). A shared k-mer immediately followed by a lower-occupancy k-mer defines a candidate left anchor, whereas a shared k-mer immediately preceded by a lower-occupancy k-mer defines a candidate right anchor. Closely spaced candidate left anchors are collapsed by retaining the one with the highest occupancy, with the leftmost retained in case of a tie. Each retained left anchor is paired with a downstream right anchor separated by at most l middle amino acids (l=35 by default); the right anchor with the highest occupancy is selected, with the closest retained in case of a tie. The complete sequence from the beginning of the left anchor to the end of the right anchor is then extracted from the original 20-amino-acid sequence. Sequences bounded by the same pair of recoded anchors form a candidate variant group.

**Figure 1:**
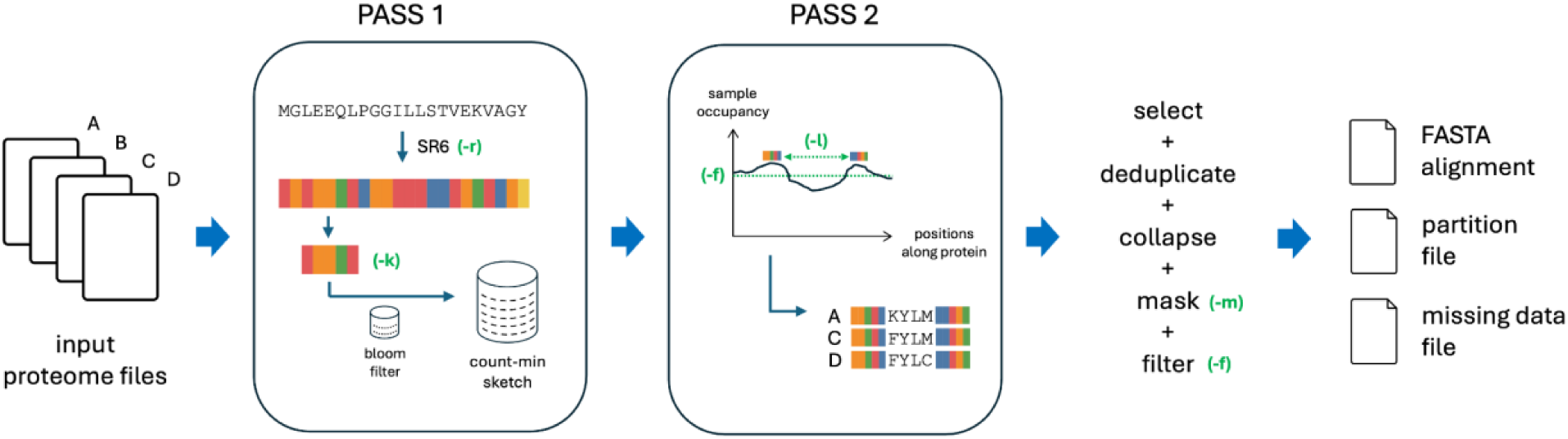
Overview of the kamino workflow. Input proteomes are processed in two passes. In the first pass, proteins are recoded into a six-state alphabet and k-mer occupancies are estimated using per-proteome Bloom filters and a count-min sketch. In the second pass, proteins are recoded again (omitted from the schematic for clarity), changes in k-mer occupancy are used to identify conserved anchors flanking variable regions, and the corresponding amino acid sequences are extracted to form candidate variant groups. Variant groups are then filtered, trimmed and concatenated to produce the final alignment and associated output files. kamino parameters are shown in green and in parentheses at the points where they apply.

Because the same pair of anchors can occasionally delimit blocks of different lengths, for example due to indels, kamino determines the number of proteomes supporting each block length. Only the uniquely best-supported length is retained, and it must occur in at least ⌈f × N⌉ proteomes; groups where two lengths are equally well supported are discarded, so that all sequences within a variant group have the same length and are pseudo-aligned. Variant groups can also overlap, since several nearby anchor pairs may describe the same protein region. To avoid counting the same sequence signal twice, groups are ranked primarily by the number of proteomes supporting them and secondarily by length. Starting from the highest-ranked group, kamino removes subsequent groups that share recoded sequence with an already retained group, detecting overlap using recoded 21-mers from the extracted blocks. The result is a non-overlapping set of variant groups preferentially supported by the largest number of proteomes. For each retained variant group, kamino reconstructs one amino acid sequence per proteome. A proteome lacking the group is represented by gaps (-). Where several copies of the same group occur within a proteome, they are collapsed into a single consensus sequence: identical amino acid states are retained and conflicting states become X. This effectively merges closely related sequences, typically in-paralogs, into one sequence per proteome.

kamino then filters the reconstructed groups in three steps. First, it builds a majority-rule consensus in the recoded alphabet; if a proteome’s sequence contains at least m consecutive differences from this consensus (m=5 by default), its complete sequence for that variant group is masked as X. This removes long runs of divergence that may reflect genuine but unwanted polymorphism, such as micro-inversions (Chaisson et al. 2006; Goldberg et al. 2014), or misalignment from multiple indels (Derelle et al. 2025). Setting m to 0 disables masking, whereas setting it to 1 results in an empty alignment.

Second, columns within the middle region are retained only when amino acids are present in at least the fraction of proteomes specified by f, with gaps and X treated as missing data (f=0.85 by default); the retained middle region is trimmed to its first and last polymorphic positions in the recoded alphabet, and groups with no polymorphic middle position are discarded. Third, up to c positions from the beginning of the right anchor are appended to the retained middle region (c=3 by default), provided they also satisfy the minimum occupancy threshold. These neighbouring positions supply constant sites for phylogenetic analyses, though positions constant in the six-state alphabet can be variable in the original amino acid alphabet.

Each variant group that passes these filters contributes one contiguous block to the final alignment, corresponding to one partition. The resulting blocks are concatenated to form the final alignment in the original 20-amino-acid alphabet. kamino outputs the alignment in FASTA format, a partition file reporting block coordinates and consensus protein names, and a per-sample missing-data file. kamino is fully deterministic so it produces reproducible outputs for a given parameter set. When -f > 0.5, the same homologous position cannot appear more than once in the final alignment by construction: any two retained variant groups must overlap in sample occupancy and therefore cannot represent disjoint encodings of the same site.

### Additional options

We implemented two additional options for user convenience of input and output. The -- nj option generates a neighbour-joining tree alongside other outputs, with all pairwise distances computed from the alignment using a single F81 correction with fixed LG stationary amino-acid frequencies (Le and Gascuel 2008). The resulting tree provides an overview of sample relationships but is not intended for detailed phylogenetic inferences. The --genomes option accepts bacterial genomes as input instead of proteomes, predicting protein-coding genes, translating them with bacterial genetic code 11, and passing the resulting sequences to kamino as input proteomes. This gene-prediction algorithm is less accurate than full gene modelling (Hyatt et al. 2010) but is faster and avoids an external dependency when only bacterial assemblies are available.

### NCBI and GTDB testing datasets

The list of eukaryotic taxa at each taxonomic rank was downloaded from NCBI taxonomy in February 2026; prokaryotic taxa were drawn from the curated GTDB database release R10-RS226 (Parks et al. 2026). For each taxon, a custom Python script retrieved genome accessions using datasets v18.19.0 (O’Leary et al. 2024), retaining only annotated genomes and excluding atypical assemblies. Proteome files were then downloaded for genomes passing a minimum predicted protein count threshold, set to 400 for prokaryotes and 2,000 for eukaryotes, to exclude incomplete proteomes. For each taxon, the script selected a fixed number of proteomes using a greedy procedure designed to maximise taxonomic diversity below the rank analysed: sampling was first balanced across child ranks and then across deeper ranks, with at most two proteomes retained per species. Taxa were discarded if fewer than the required number of proteomes remained after filtering, or if a single child taxon represented more than 75% of the selected proteomes (e.g., the most abundant class of a given phylum should not exceed 75% of the dataset), to avoid datasets dominated by a single lower-rank group.

### Analyses of the MTBC dataset

The MTBC dataset comprised 730 isolates listed in the file out_fastlin_v3.txt from the repository https://github.com/rderelle/barcodes-fastlin. The isolates had previously been typed using fastlin v0.4.2.1 (Derelle et al. 2023) with a modified set of barcodes from TB-Profiler (Verboven et al. 2022). For these analyses, genome assemblies corresponding to the isolates were downloaded from the AllTheBacteria project (Hunt et al. 2026) and, when unavailable there, from NCBI. Genome assemblies were analysed using Mashtree v1.4.6 (Katz et al. 2019) and SKA v0.5.1 (Derelle et al. 2024), using the command line described at https://bacpop.org/guides/snp_alignment_with_ska/. The neighbour-joining tree based on the SKA alignment was inferred using RapidNJ v2.3.3 (Simonsen and Pedersen 2011). Gene prediction was performed using Prodigal v2.6.3 (Hyatt et al. 2010) to generate protein sets for the kamino analysis. The genome and proteome datasets are available in the Zenodo archive.

### Analyses of GTDB taxa

To build datasets for the phylogenetic analyses of GTDB taxa, we modified the script described above to accept a minimum and maximum number of proteomes rather than a fixed number, set to 40 and 200, respectively. For the monophyly testing, taxa represented by fewer than three isolates were excluded, compared with fewer than two isolates for the MTBC analyses, because up to two isolates per species were retained in the GTDB datasets.

### Orthogroup comparative analyses

The three sets of 40 proteomes were each clustered into orthogroups using OrthoFinder v3.1.5 (Emms et al. 2025) with the ‘-og’ option to avoid the computationally costly alignment and phylogenetic analysis of orthogroups. For this comparison, we used a modified version of kamino in which all positions from variant groups passing the standard filters were retained, ensuring that partition sequences remained contiguous subsequences of the original proteins and could be matched directly to OrthoFinder orthogroups. We also set -c 13, thereby increasing the minimum partition length from 4 to 14 amino acids, in order to facilitate exact sequence matching. These two changes did not affect the identification or selection of partitions, only the number of positions retained within each partition. Partition sequences containing - or X were discarded, and the remaining sequences were matched exactly to proteins and assigned to their OrthoFinder orthogroups using a custom Python script. Partitions with fewer than 30 sequences assigned to orthogroups were excluded. This script, the modified version of kamino, and all output files are available in the Zenodo archive. The Drosophila phylogeny was inferred using the alignment generated by the unmodified version of kamino with default parameters.

### BUSCO analyses

BUSCO marker sets version odb12 were downloaded from https://busco-data.ezlab.org/v5/data/lineages/; marker counts per set are provided in Supplementary Data. Markers were identified in proteome files using BUSCO v6.0.0 (Manni et al. 2021) in protein mode. Identified markers were combined into concatenated phylogenomic alignments using BUSCO_phylogenomics (https://github.com/jamiemcg/BUSCO_phylogenomics), with gene-tree building disabled and a minimum single-copy gene threshold of 85%. Both tools were run using 4 CPUs, and reported runtimes for BUSCO analyses correspond to their combined wall-clock times.

### Mammalian dataset

For the mammalian analyses, we used the 190 species included in the OrthoMaM v12a database (Allio et al. 2024). Predicted proteomes from the corresponding RefSeq genome assemblies were downloaded from NCBI using NCBI Datasets. The proteome dataset is available in the Zenodo archive.

### Quality assessments and phylogenetic analyses

Alignment difficulty scores were inferred using Pythia v2.0.0 (Haag et al. 2022) with RAxML-NG v2.0.0-beta3 (Kozlov et al. 2019) under default parameters. Saturation levels were computed using the saturation command in PhyKIT v2.1.90 (Steenwyk et al. 2021), applied to alignments and their corresponding IQ-TREE trees. Phylogenetic analyses were run using IQ-TREE v3.0.1 (Wong et al. 2025) under the LG+I+G model and using FastTree v2.2.0 double-precision (Price et al. 2010) under the LG model. BioNJ trees were retrieved from IQ-TREE output. For the mammalian dataset, an additional IQ-TREE analysis was performed using the PMSF approximation to the site-heterogeneous LG+C20+I+G model, with the LG+I+G tree used as the guide tree. Cluster information distances and Robinson-Foulds distances between unrooted trees were calculated using the R package TreeDist v2.12.0 (Smith 2022) and used to build neighbour-joining trees with the R package ape v5.8.1 (Paradis et al. 2004) that were rooted using the R package phangorn v2.12.1 (Schliep 2011). Runtime and memory measurements were conducted on an Intel Xeon Platinum 8358 CPU @ 2.60 GHz.

### Code and data access

The kamino codebase is available at https://github.com/rderelle/kamino and precompiled binaries can be installed via Bioconda. Analyses were performed with kamino v2.1.1 using default parameters unless specified otherwise. All alignments, phylogenetic trees, and Python and R scripts used in this study have been deposited in Zenodo: https://zenodo.org/records/20294318.

## Results

### kamino performance declines at high divergence

The requirement for conserved sequences to be shared across most proteomes is expected to yield less signal as diversity increases, as with other reference-free methods (Derelle et al. 2024, 2025). To quantify this effect, we built amino acid phylogenomic alignments from 40 proteomes sampled across increasing taxonomic ranks in prokaryotes and eukaryotes (see Materials and methods), comprising 93 eukaryotic taxa and 269 prokaryotic taxa. Alignment size decreased slightly from genus to phylum in eukaryotes, but much more markedly in prokaryotes (Figure 2; values in Supplementary Data). Taxa with secondarily reduced proteomes generally produced smaller alignments, including Microsporidia and Mycoplasmopsis. Constant sites represented on average 48% of alignment positions (Supplementary Figure S1), while missing data increased from roughly 5% at the genus level to 10% at the phylum level (Supplementary Figure S2).

**Figure 2:**
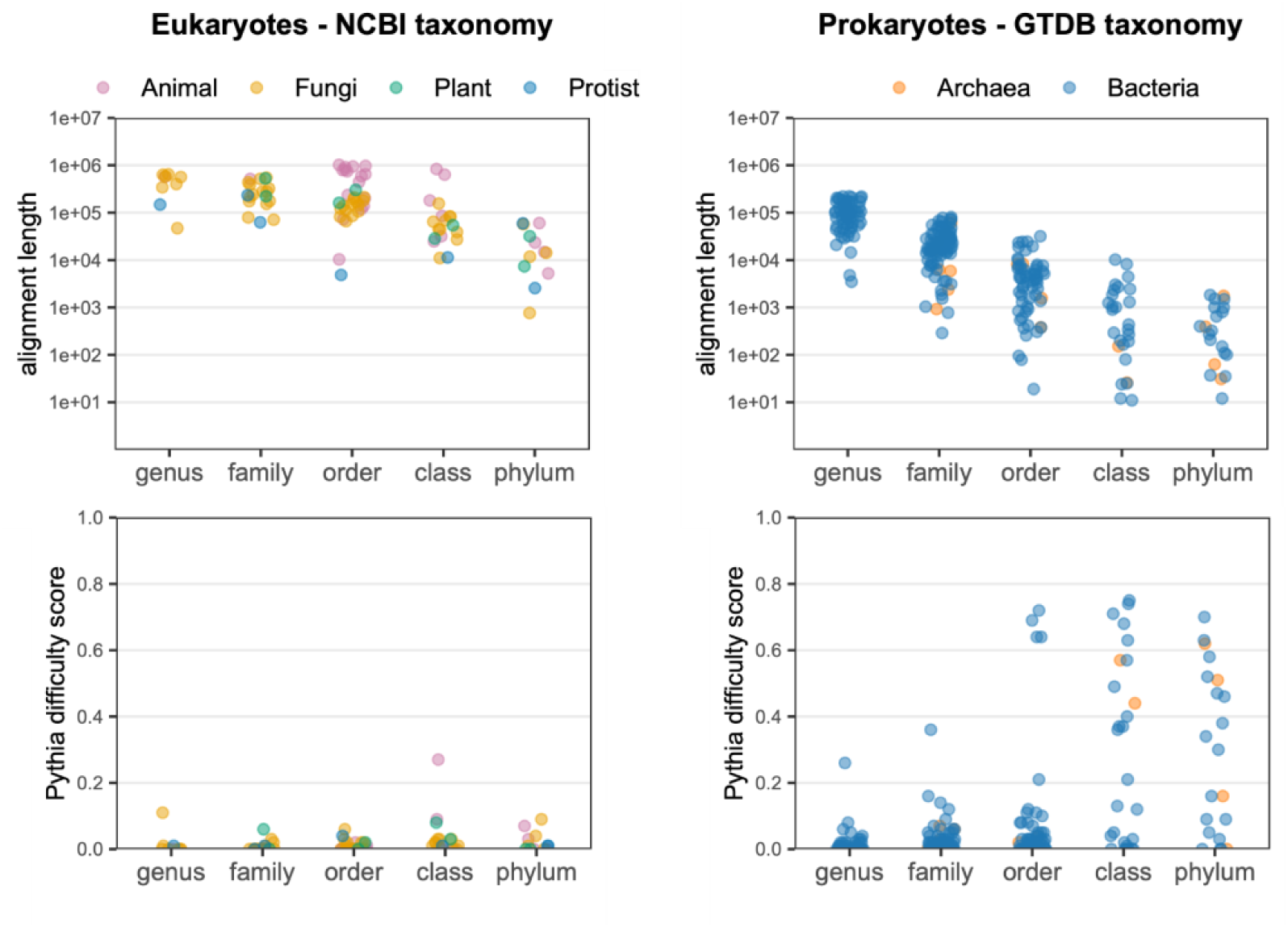
kamino analyses of 40-proteome datasets. Top panels show alignment lengths at different taxonomic ranks for eukaryotic datasets (NCBI taxonomy, left) and prokaryotic datasets (GTDB taxonomy, right). Bottom panels show Pythia difficulty scores for the same datasets (lower scores indicate easier phylogenetic inference). "Protist" refers to eukaryotic taxa outside animals, fungi, and plants. Note that the y-axes in the top panels are log-scaled, and that taxonomic groups are naturally nested, leading to some isolates occurring in more than one dataset.

We then assessed phylogenetic signals using the Pythia difficulty score (0–1), for which lower scores indicate stronger signal (Haag et al. 2022). Eukaryotic alignments produced scores below 0.1, with the exception of the class Aves (animal) whose deep relationships are affected by rapid radiations and limited phylogenetic signal (Zhang and Ma 2024; Suh et al. 2015). Prokaryotic datasets produced low scores at the genus and family ranks, but scores up to 0.8 at higher ranks. However, alignments with scores above 0.2 were consistently the shortest (Supplementary Figure S3), suggesting these reflected insufficient data rather than noisy or conflicting signals.

Users can increase kamino’s sensitivity by relaxing the minimum sample occupancy threshold (-f parameter). Lowering it from 0.85 to 0.75 increased alignment length by roughly 1.5- to 2-fold in most datasets, with the largest gains in the shortest alignments (Supplementary Figure S4). Even so, the same taxonomic limitations persisted.

### kamino partitions largely match OrthoFinder orthogroups

To assess whether kamino partitions correspond to coherent orthologous protein groups, we matched kamino partition sequences to orthogroups independently inferred by OrthoFinder (see Materials and methods). We performed this analysis on three genus-level datasets of 40 proteomes each: Achromobacter, Drosophila, and Aspergillus. We found that 98.1%, 84.9%, and 93.5% of kamino partitions had all sequences assigned to the same orthogroup in Achromobacter, Drosophila, and Aspergillus, respectively (Figure 3A). When the comparison was restricted to orthogroups present in at least 30 of the 40 proteomes, these percentages increased to 99.9%, 94.2%, and 97.2%, respectively. These results show that discordance between kamino partitions and OrthoFinder orthogroups was substantially associated with low-occupancy orthogroups, suggesting that mixed assignments may reflect over-splitting by OrthoFinder rather than distinct, broadly distributed paralogous groups.

**Figure 3:**
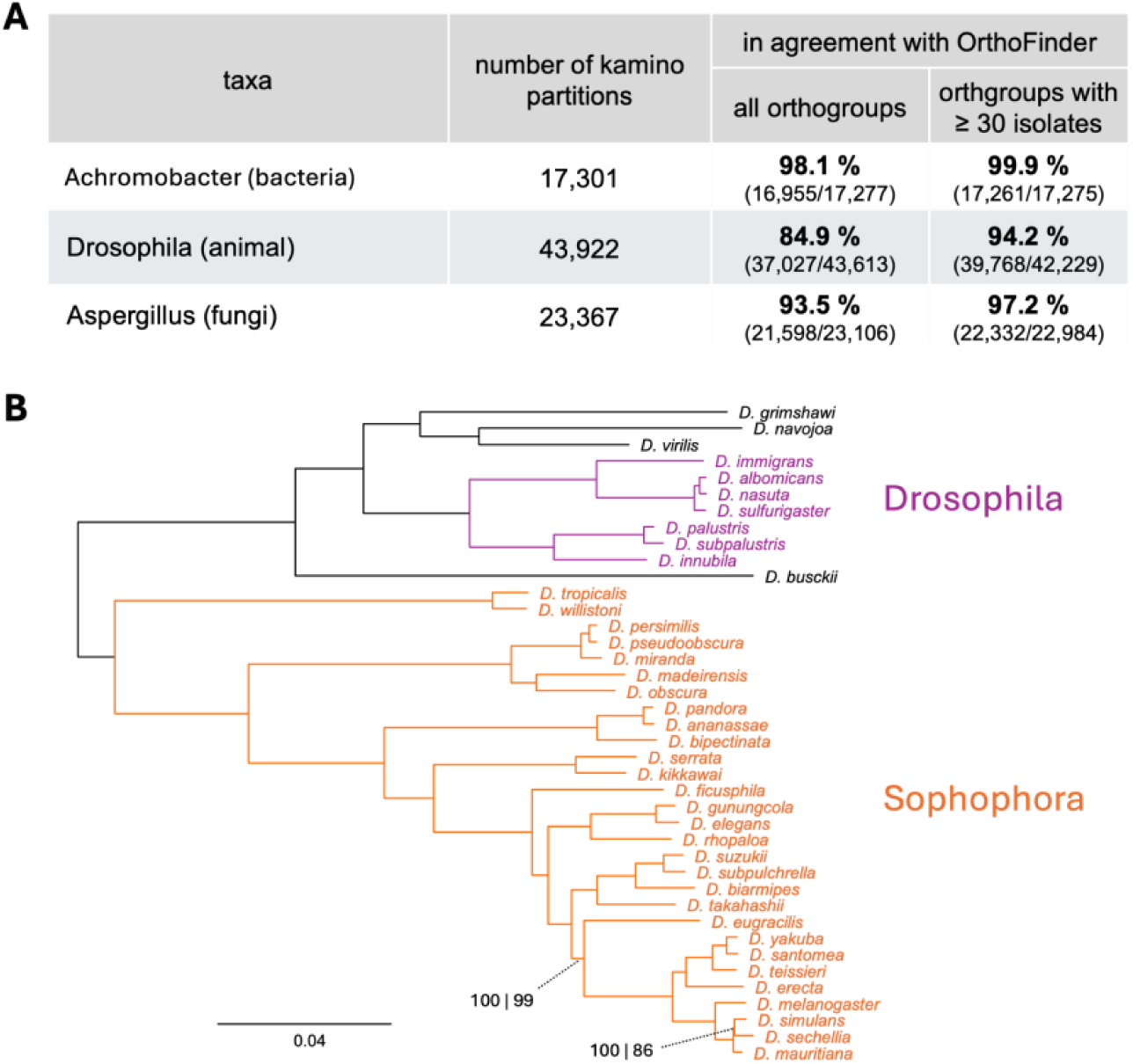
Comparison with OrthoFinder orthogroups. **A**. Agreement between kamino partitions and OrthoFinder orthogroups across the three 40-proteome datasets. Values in parentheses indicate the number of agreeing partitions over the number compared. **B**. IQ-TREE phylogeny of the Drosophila dataset inferred from kamino partitions agreeing with OrthoFinder, using LG+I+G and 2,000 ultrafast bootstrap replicates. The same topology was recovered from disagreeing partitions. Bootstrap values are shown only when below 100% in at least one analysis, with values given for agreeing | disagreeing partitions. The two main subgenera represented in the dataset, Sophophora and Drosophila, are coloured for clarity.

We then performed phylogenetic analyses on the Drosophila dataset, which showed the lowest agreement between kamino partitions and OrthoFinder orthogroups. We analysed separately partitions in full agreement with OrthoFinder orthogroups (490,569 positions from 37,027 partitions) and partitions showing disagreement (77,203 positions from 6,586 partitions). The two independent alignments yielded the same topology (Figure 3B), consistent with the known relationships among these Drosophila species (Finet et al. 2021; Suvorov et al. 2022). The tree inferred from partitions in full agreement with OrthoFinder had 100% bootstrap support for all branches, whereas the tree inferred from discordant partitions had lower support at two branches. These differences may simply reflect the smaller size of the latter alignment, as bootstrap support tends to increase with dataset size (Minh et al. 2020; Salichos and Rokas 2013). Overall, the two sets of partitions contained similar phylogenetic signals, indicating that kamino partitions showing discordance with OrthoFinder orthogroups still retained meaningful phylogenetic information.

### kamino recovers curated bacterial classifications

We evaluated kamino’s ability to recover curated classifications across different evolutionary scales. The first benchmark used 730 isolates from the Mycobacterium tuberculosis complex (MTBC), for which fine-scale lineages had previously been defined phylogenetically from variants called by read alignment (Napier et al. 2020). The dataset contained 136 lineages, 129 of which were represented by at least two isolates. We compared kamino with the sketch-based Mashtree and the split k-mer-based SKA, both of which are alignment-free. kamino recovered the MTBC lineage structure as accurately as SKA, which has previously been validated for this task (Chilengue et al. 2026): only 3 of the 129 lineages were non-monophyletic in the FastTree analysis, the same three as with SKA (Figure 4A; non-monophyletic lineages listed in Supplementary Data). kamino also showed high efficiency with runtimes comparable to that of Mashtree. Finally, running kamino directly from genome assemblies yielded a shorter alignment due to the rapid, approximate gene prediction step (135,541 versus 143,880 positions), but produced nearly identical phylogenetic results to the default proteome-based analysis.

**Figure 4:**
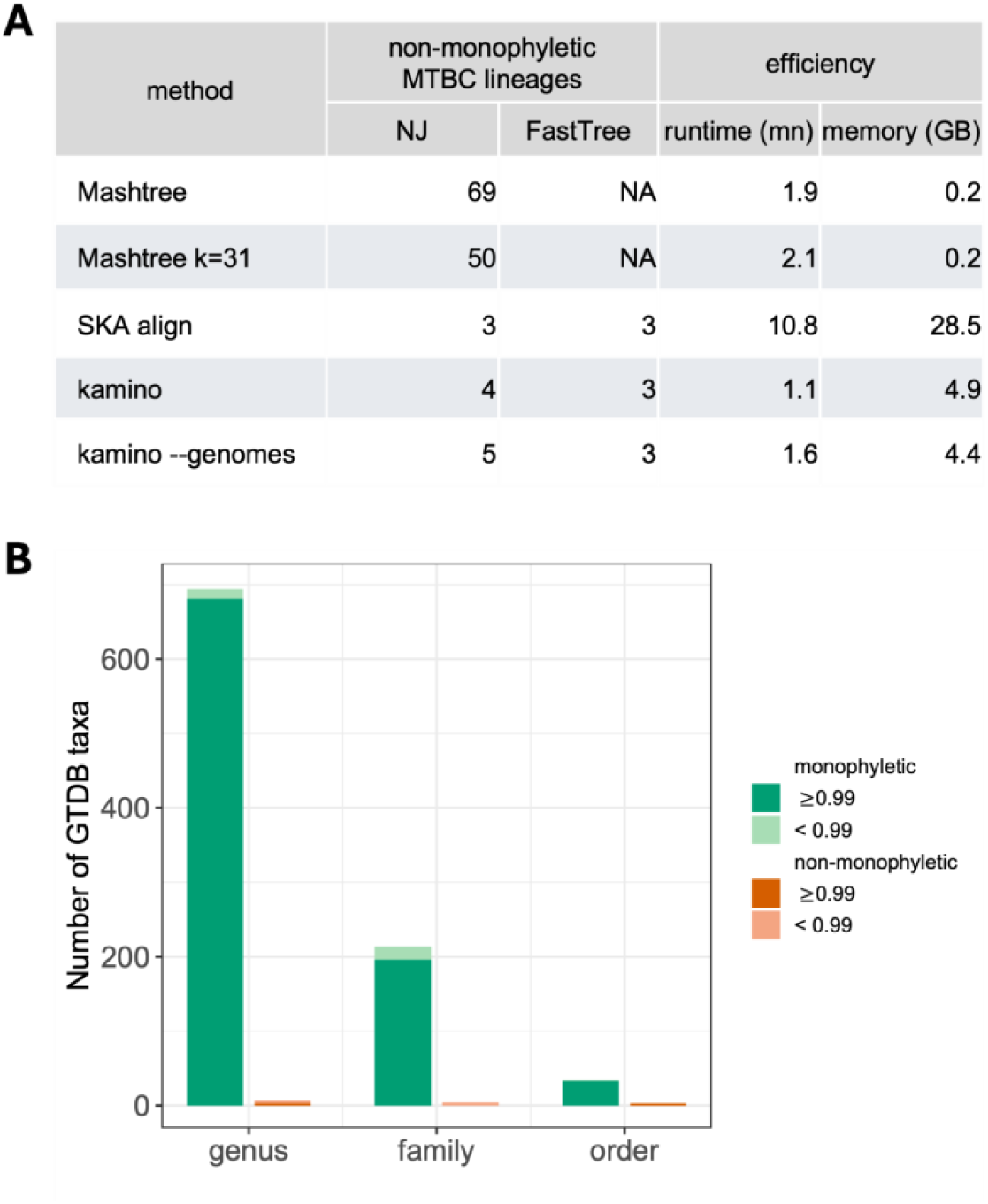
kamino benchmarking against curated classifications. **A**. Analysis of a dataset of 730 MTBC isolates, containing 129 lineages with at least two isolates. The table shows the numbers of non-monophyletic lineages (the lower the better) and efficiency metrics obtained by the different methods, all run using 4 CPUs. Default parameters were used unless specified: ‘Mashtree k=31’ indicates an increase of the k-mer size from 21 (default) to 31, and ‘kamino –genomes’ indicates a kamino run using genome assemblies as input. NJ: neighbour-joining; NA: non-applicable; mn: minutes. **B**. Analysis of GTDB taxa. The bar plot reports the numbers of monophyletic and non-monophyletic GTDB taxa in kamino-derived FastTree analyses under the LG model and with SH-like branch support values.

At a broader evolutionary scale, we compared kamino phylogenies with the GTDB taxonomy. We regenerated kamino alignments for the GTDB taxa analysed above, this time allowing up to 200 isolates per dataset rather than a fixed set of 40. Alignments shorter than 2,000 positions were excluded, and phylogenetic trees were inferred with FastTree. For each tree, we tested monophyly of taxa at the immediately lower taxonomic rank represented by at least three isolates (e.g., genera within family-level datasets), yielding 125 phylogenetic trees and tests of 956 GTDB taxa spanning genera to orders. kamino showed strong agreement with the curated taxonomy: 942 of 956 taxa were found monophyletic, 910 of these with SH-like branch support ≥0.99, while only 6 were non-monophyletic with support ≥0.99 (Figure 4B; non-monophyletic taxa listed in Supplementary Data); all six corresponded to paraphyly rather than polyphyly.

### kamino and BUSCO show broadly similar phylogenetic signals

We compared alignments generated by kamino with those produced by a BUSCO marker-based pipeline across four taxa, each represented by 120 proteomes: the bacterial genus Mycobacterium, the bacterial family Enterobacteriaceae, the fungal order Hypocreales, and the animal class Insecta. For each dataset, we used both a broad BUSCO marker set and a larger taxon-specific marker set, hereafter referred to as “BUSCO broad” and “BUSCO taxa”, respectively. With the exception of Insecta, kamino alignment lengths were similar to those obtained with BUSCO broad, whereas BUSCO taxa produced substantially larger alignments (Figure 5). Pythia difficulty scores were low and similar across all datasets, ranging from 0 to 0.06. kamino alignments were also consistently less saturated than BUSCO-based alignments, with the largest differences observed in the eukaryotic datasets.

**Figure 5:**
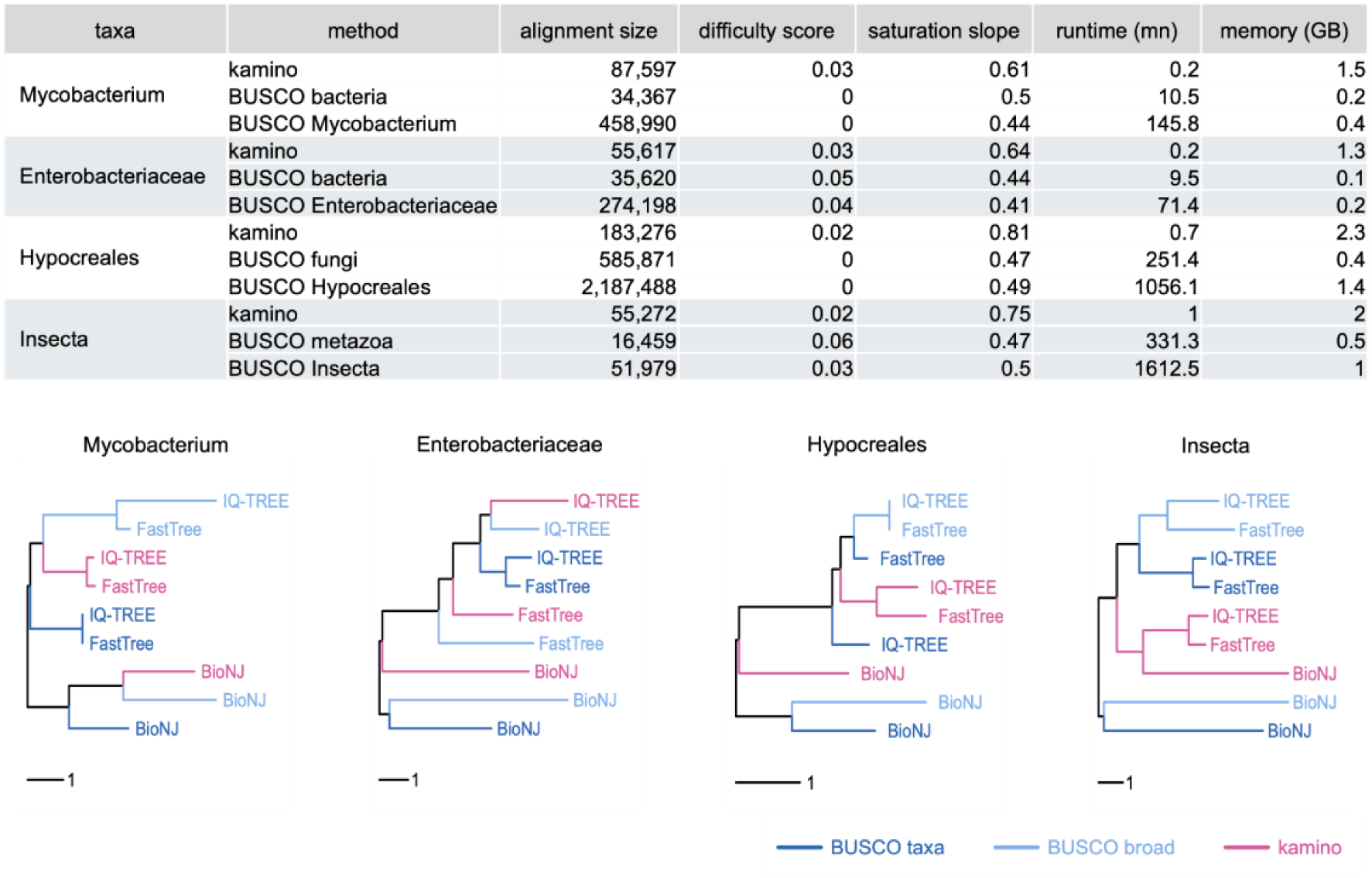
Comparative analyses of kamino and a BUSCO-based pipeline. The top table summarises, for each dataset and alignment method, the alignment length, Pythia difficulty score, saturation slope, runtime, and memory usage using 4 CPUs. The lower panels show neighbour-joining trees of trees built from pairwise clustering information distances among the nine inferred phylogenies for each dataset: three alignment strategies, kamino, BUSCO broad, and BUSCO taxa, each analysed with BioNJ, FastTree, and IQ-TREE. Trees are midpoint rooted. Tip labels indicate the inference method, and colours indicate the alignment source. Trees based on Robinson-Foulds distances are provided in Supplementary figure S5.

We next compared the phylogenetic signals recovered from each alignment using BioNJ, pseudo-maximum likelihood with FastTree, and maximum likelihood with IQ-TREE. For each dataset, pairwise distances among the nine resulting trees were used to construct a neighbour-joining tree summarising their relationships (Figure 5). For Mycobacterium, Enterobacteriaceae, and Hypocreales, trees grouped mainly by inference method rather than alignment source, indicating that kamino and BUSCO alignments contained broadly similar, although not identical, phylogenetic signals. In contrast, the three kamino trees clustered together for Insecta, indicating a stronger effect of alignment source in this dataset. Examination of deep relationships showed that only the kamino trees recovered Condylognatha (Thysanoptera + Hemiptera), whereas only the BUSCO-based trees recovered Palaeoptera (Odonata + Ephemeroptera) (Supplementary Figures S6 and S7). The latter relationship remains debated (Simon et al. 2018).

### kamino reproduces major mammalian relationships

We finally examined phylogenetic relationships within mammals, for which several deep relationships have historically been difficult to resolve (Foley et al. 2016; Esselstyn et al. 2017). We used the dataset of 190 species from the OrthoMaM v12a database (Allio et al. 2024), and only performed kamino analyses, as BUSCO analyses would be computationally demanding for a dataset composed of such large proteomes. Using default parameters, kamino generated an alignment of 1,243,961 positions in 4 minutes using 22.4 GB of memory on four CPUs. To reduce the computational cost of downstream phylogenetic analyses, we increased the minimum sample occupancy from 0.85 to 0.95 (-f parameter), yielding a shorter alignment of 576,374 positions in 3.3 minutes using 15.2 GB of memory.

Phylogenetic analyses were performed on this alignment under the LG and site-heterogeneous LG+C20-PMSF models, with 2,000 ultrafast bootstrap replicates, and both models yielded the same tree topology. Under the LG model, several deep branches received relatively low bootstrap support, including the root of Placentalia separating Xenarthra+Afrotheria from Euchontoglires+Laurasiatheria and the position of Scandentia within Euarchontoglires, two relationships that remain contentious (Figure 6; the full trees are available in Supplementary Figures S8 and S9). In contrast, all branches received 100% bootstrap support under the site-heterogeneous LG+C20-PMSF model. This fully supported deep topology was largely congruent with the whole-genome phylogeny of 241 placental mammals reported by (Foley et al. 2023), although kamino recovered Perissodactyla as sister to Cetartiodactyla (Euungulata) whereas Foley et al. recovered it as sister to Ferae (Zooamata). Notably, the kamino alignment was generated directly from proteomes in only a few minutes, whereas comparable mammalian phylogenomic analyses rely on computationally intensive whole-genome alignments.

**Figure 6:**
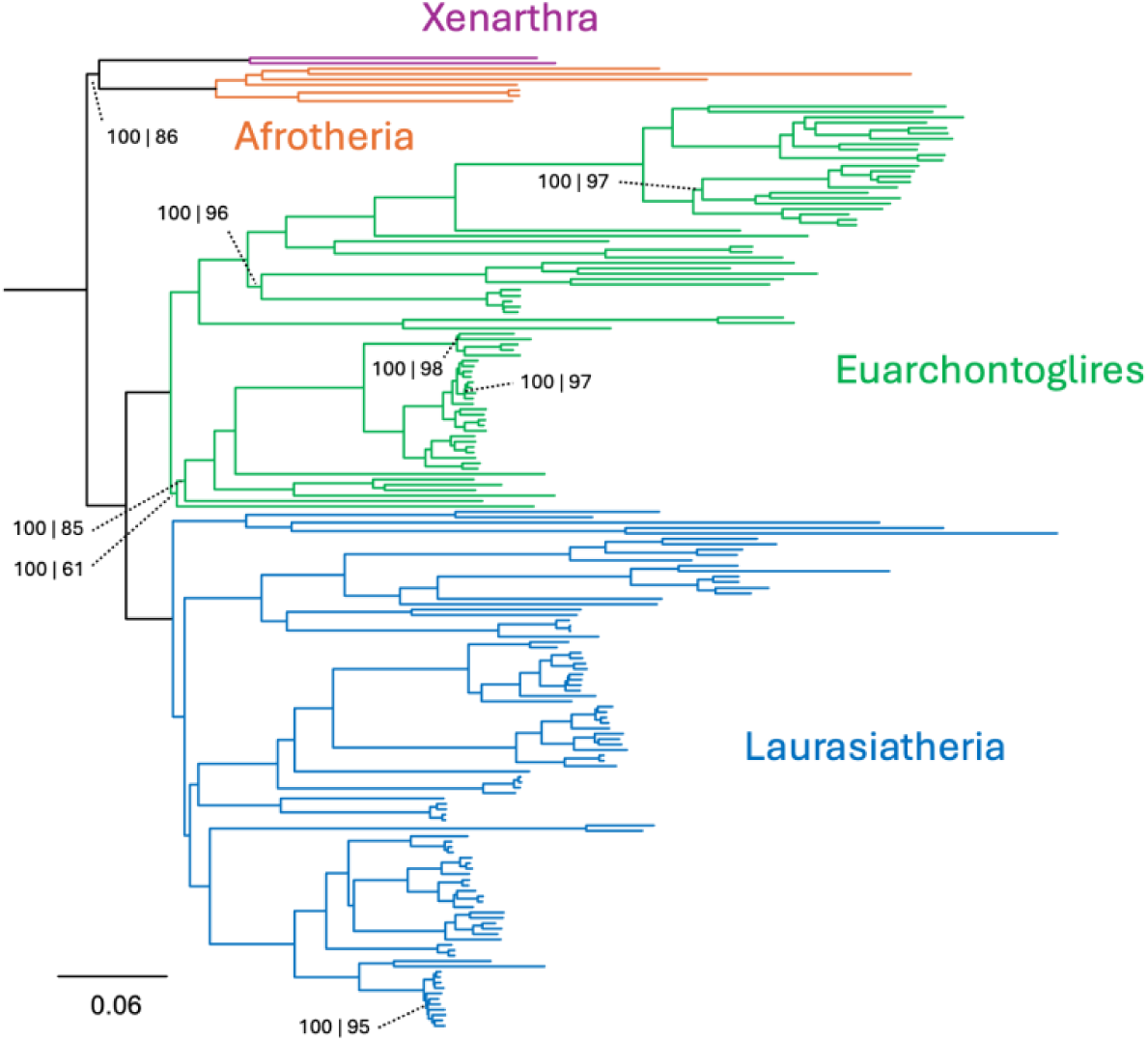
Phylogenetic analyses of 190 mammals. IQ-TREE phylogeny of the OrthoMaM v12a dataset inferred under the LG+C20-PMSF+I+G model with 2,000 ultrafast bootstrap replicates. The 10 non-placental mammals used to root the tree were removed for clarity. The same topology was recovered under the LG+I+G model. Bootstrap values are shown only for branches with support below 100% in at least one analysis, with values given as LG+C20-PMSF | LG.

### kamino is fast and scales approximately linearly

In the MTBC analyses, kamino showed efficiency comparable to that of the fast genome-based programs SKA and Mashtree. Compared with the BUSCO marker-based pipeline across the four 120-proteome datasets, kamino was one to two orders of magnitude faster for the prokaryotic datasets and two to three orders of magnitude faster for the eukaryotic datasets, depending on the size of the BUSCO marker set used (Figure 5).

We next examined how runtime and memory use scaled with dataset size by rerunning kamino on each of the four taxa with 40, 80, 120, 160, and 200 proteomes (4 CPUs throughout). Both runtime and memory use increased approximately linearly with the number of proteomes, with steeper increases for eukaryotic datasets, which have larger proteomes (Figure 7). Because kamino does not retain the full k-mer diversity in memory, memory use is driven mainly by the sequences stored in variant groups. Memory requirements can therefore become substantial for datasets with large proteomes that yield many variant groups, as illustrated by the mammalian analyses above. In such cases, users can increase kamino’s minimum sample occupancy threshold, thereby reducing the number of variant groups detected and lowering memory use. Overall, kamino runtimes and memory requirements should allow rapid analyses of datasets comprising a few thousand prokaryotic genomes or several hundred eukaryotic genomes.

**Figure 7:**
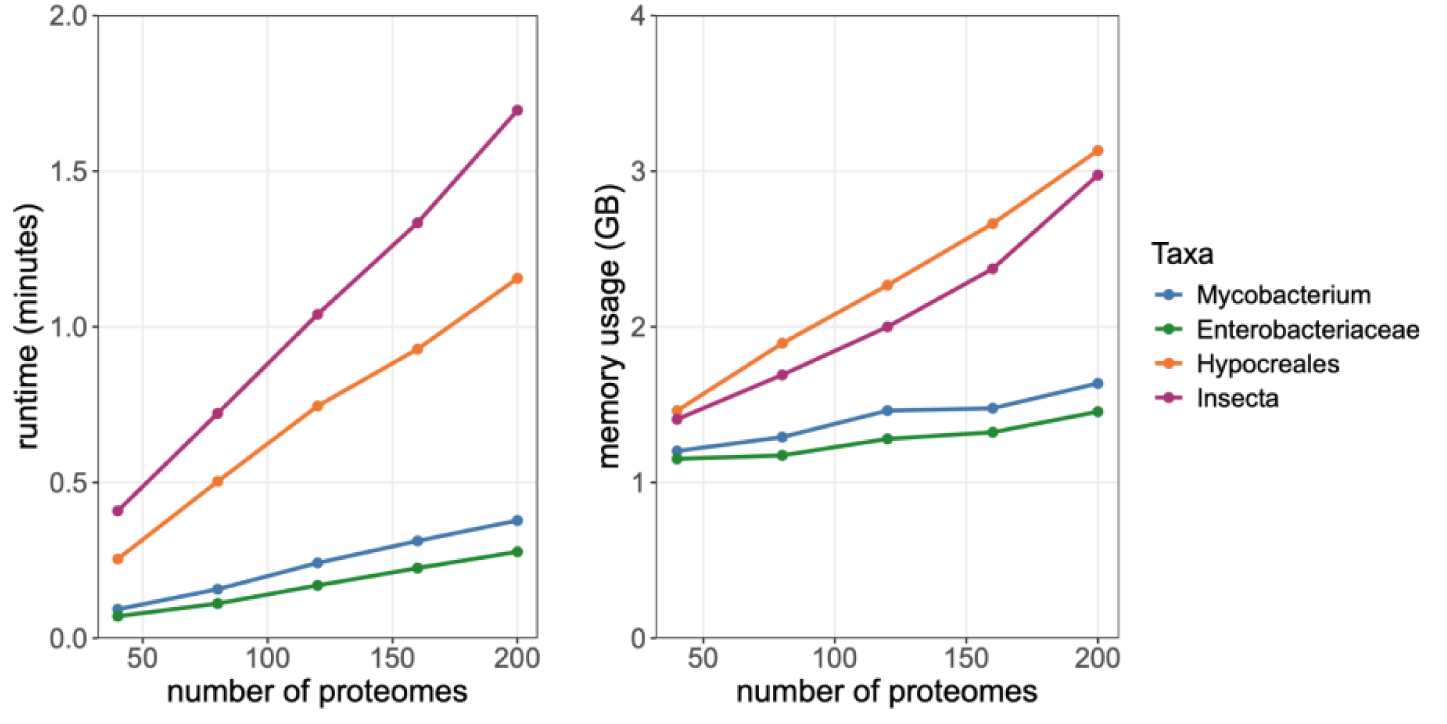
kamino efficiency testing. The two panels show kamino runtime and maximum memory use for increasing dataset sizes using 4 CPU threads. Numbers of positions in kamino alignments are available in Supplementary Figure S10.

Finally, we evaluated multithreading by reanalysing the 120-proteome datasets using 1 to 12 CPUs. Runtime decreased markedly from 1 to 4 CPUs, with more modest gains thereafter (Supplementary Figure S11). This suggests that using many additional CPUs provides limited further speed-up for kamino, and that comparisons with BUSCO-based pipelines should be interpreted with some caution, as all-versus-one approaches can be more easily parallelised and may therefore benefit more from additional CPUs.

## Discussion

Amino acid-based phylogenetics has largely relied on the same workflow: identifying orthologous proteins, aligning them, removing paralogs, and concatenating the resulting alignments. This approach has remained the standard because it is highly effective: it can produce alignments across most taxonomic scales and can be made scalable. Its main drawback is speed, as the multiple alignment steps slow analyses and require pipelines involving several programs. kamino was developed to remove this bottleneck. By identifying conserved k-mer anchors flanking short variable regions across proteomes, it builds phylogenomic alignments directly, without orthogroup inference or sequence alignment, making it faster and simpler to run than classical marker-based workflows. The OrthoFinder comparison supports this strategy: despite making no explicit orthology assignments, most kamino partitions grouped sequences from the same orthogroup. The similar phylogenetic signal recovered from discordant partitions also highlights that orthogroup inference is itself an approximation. Gene duplication and loss, but also fusion, fission and domain rearrangement, can give different regions of the same protein distinct evolutionary histories, making whole-protein orthology assignments ambiguous (Sjölander et al. 2011; Gabaldón and Koonin 2013). Errors in gene prediction compound this problem, particularly in eukaryotes, where split or merged gene models can substantially affect inferred orthology relationships (Prieto-Baños et al. 2025).

kamino alignments retained substantial phylogenetic signal across very different evolutionary depths. They recovered MTBC lineages as accurately as SKA, agreed closely with curated GTDB taxa, recovered the expected relationships among the Drosophila species analysed, and produced mammalian relationships largely congruent with recent whole-genome phylogenies. Comparisons with BUSCO alignments revealed broadly similar phylogenetic signals in three of the four datasets, while kamino alignments were consistently less saturated across all four. The strong performance of kamino on mammals is consistent with the success of ultraconserved elements flanked by more variable regions in resolving difficult relationships within this group (Esselstyn et al. 2017; McCormack et al. 2012). kamino relies on a conceptually similar strategy at the amino-acid level, using conserved flanking sequences to identify short variable regions. Sampling many such regions across the proteome without requiring a reference or conventional multiple-sequence alignment can therefore provide sufficient phylogenetic information for both shallow and relatively deep relationships, while substantially reducing the computational cost of alignment construction.

Given its design, kamino is less suitable at the two extremes of the genomic diversity spectrum. At very low divergence, within individual lineages such as pathogen outbreaks, the main objective is typically to recover as many variants as possible. In this setting, amino acid recoding into a six-letter alphabet will discard some informative variation, and the absence of genomic coordinates prevents the detection and removal of recombination tracts (Posada and Crandall 2002; Didelot and Wilson 2015). Genome-based alignment-free methods are therefore more appropriate for these low-divergence settings. At the opposite extreme, very high evolutionary divergence reduces the number of conserved sequences shared across proteomes and therefore limits the identification of variant groups. In the datasets tested here, kamino generated useful alignments between lineages and remained effective at genus and family levels in prokaryotes and up to the phylum level in eukaryotes, but performance declined for more divergent groups, particularly in prokaryotes. This probably reflects the higher evolutionary divergence and greater gene-content turnover found across broad prokaryotic groups (McInerney et al. 2017; Molari et al. 2025). When this occurs, the variant-extraction process yields shorter alignments rather than a noisy signal. Users can therefore simply assess whether the resulting alignment is sufficiently long, tune kamino’s parameters to recover a longer alignment, or switch to a traditional orthology- and alignment-based approach.

A separate limitation concerns the partitioning of kamino alignments. Individual partitions are short (4 to 38 positions by default), which rules out analyses that need larger, gene-level units: they are too short to infer meaningful gene trees, identify phylogenomic outliers, or examine gene-wise phylogenetic conflict, and they are not suitable as independent partitions for fitting separate substitution models. Like most marker-based pipelines, kamino does not currently perform explicit phylogenomic outlier detection, so it can potentially be affected by hidden paralogy, incomplete lineage sorting, and lateral gene transfer (Steenwyk et al. 2023; Comte et al. 2023). Future developments could address this limitation by mapping variant groups back to the proteins from which their amino acid sequences were extracted and combining them into larger orthogroup-like units, while maintaining an alignment- and reference-free approach. Such units could support outlier detection, phylogenetic analyses using separate substitution models, gene-wise concordance analyses (Minh et al. 2020), and gene-tree-based species-tree inference (Mirarab et al. 2014).

Given its speed and simplicity, kamino is primarily designed to build phylogenetic alignments quickly, on modest hardware, and without requiring prior expertise in phylogenomic workflows. Within its range of applicability, kamino provides a practical complement to, or substitute for, classical alignment-based approaches.

## Acknowledgments

The authors wish to thank members of the microbioinfo Slack channel for discussions that initiated the development of kamino, and Torsten Seemann for helpful suggestions. R.D. and L.C. acknowledge funding from the MRC Centre for Global Infectious Disease Analysis (reference MR/X020258/1), funded by the UK Medical Research Council (MRC). This UK funded award is carried out in the frame of the Global Health EDCTP3 Joint Undertaking. J.A.L. was supported by the European Molecular Biology Laboratory, European Bioinformatics Institute.

