## Supplementary figures for "kamino: fast proteome-wide variant calling for amino acid phylogenomics"

**Figure S1**
Percentages of constant positions in kamino alignments. The datasets plotted here correspond to those analysed in Figure 2. Values are available in Supplementary data.

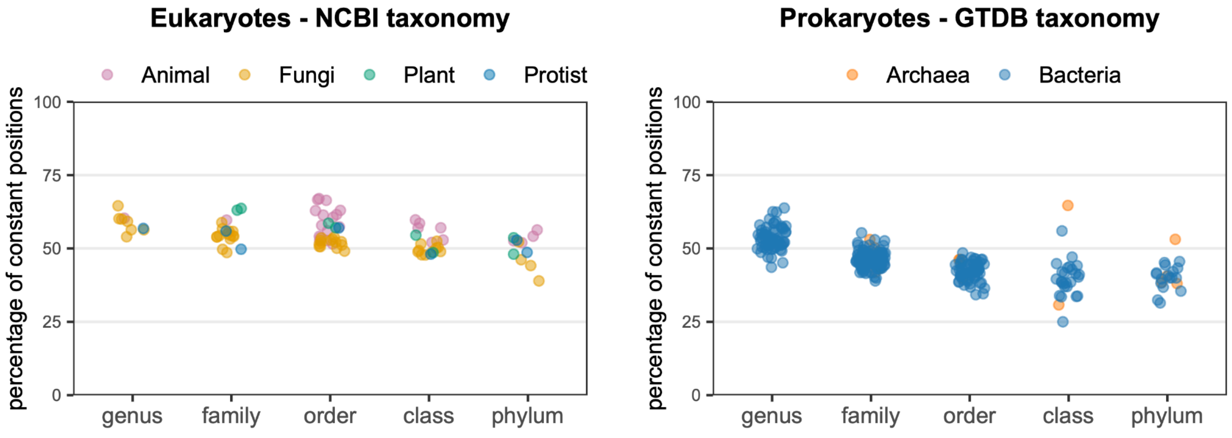


**Figure S2**
Percentages of missing data in kamino alignments. The datasets plotted here correspond to those analysed in Figure 2. Values are available in Supplementary data.

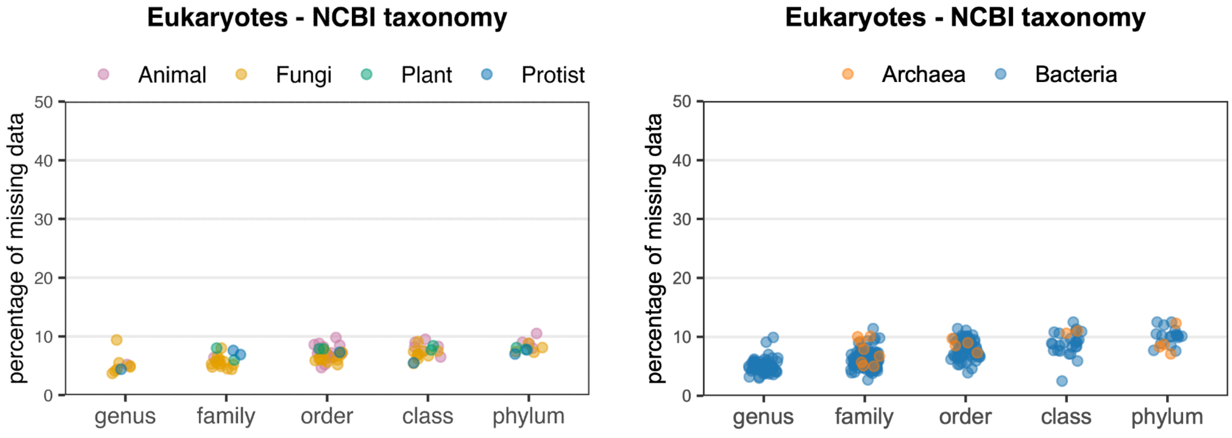

**Figure S3**
Pythia difficulty score as a function of alignment length for the 40-isolates eukaryotic and prokaryotic datasets. Please note the log scale on the x-axis.

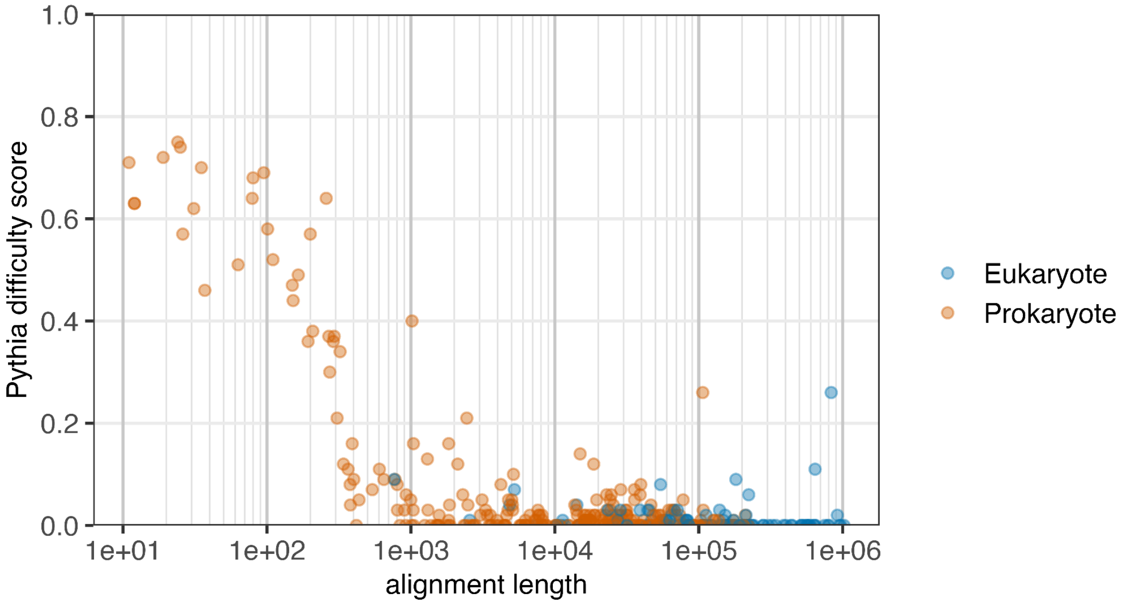


**Figure S4**
Comparison of alignment lengths obtained with default parameters and with -f 0.75 for the 40-proteome datasets. The dashed line indicates equal alignment length between the two settings.

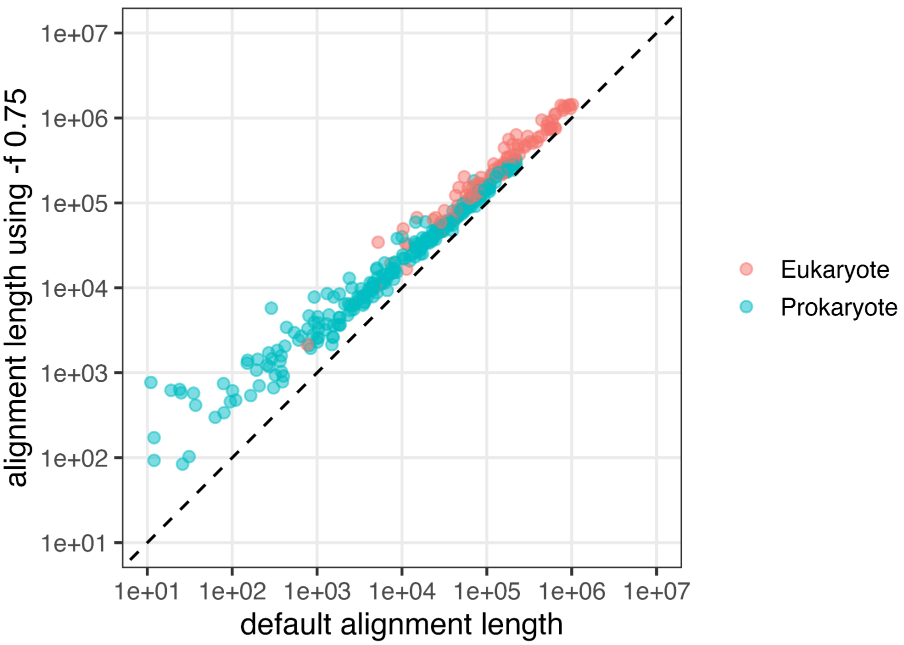

**Figure S5**

Trees of trees from the 120-proteome datasets based on Robinson-Foulds distances. Trees are midpoint rooted. Tip labels indicate the inference method, and colours indicate the alignment source.

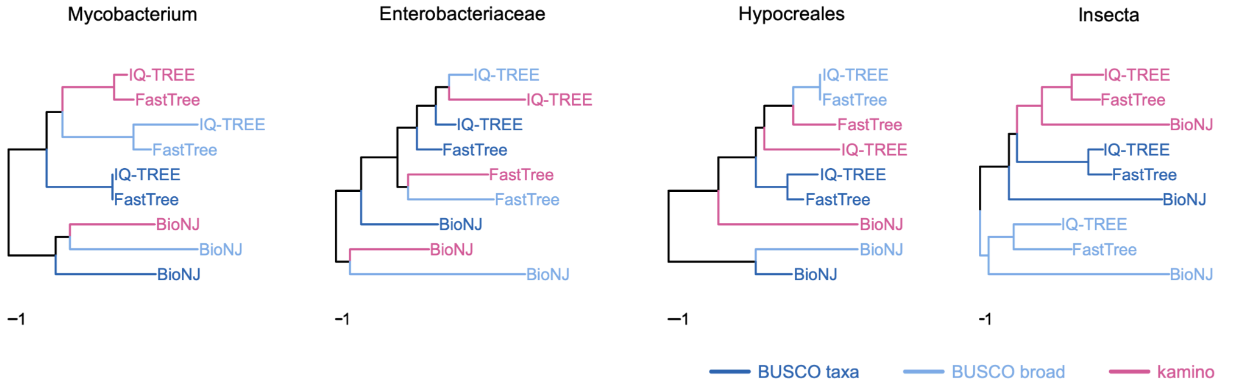


**Figure S6**

Phylogenetic tree inferred with IQ-TREE from the kamino alignment of class Insecta. The tree was rooted using Thermobia domestica (Zygentoma) as the outgroup, and order names are shown for orders represented by at least two samples.

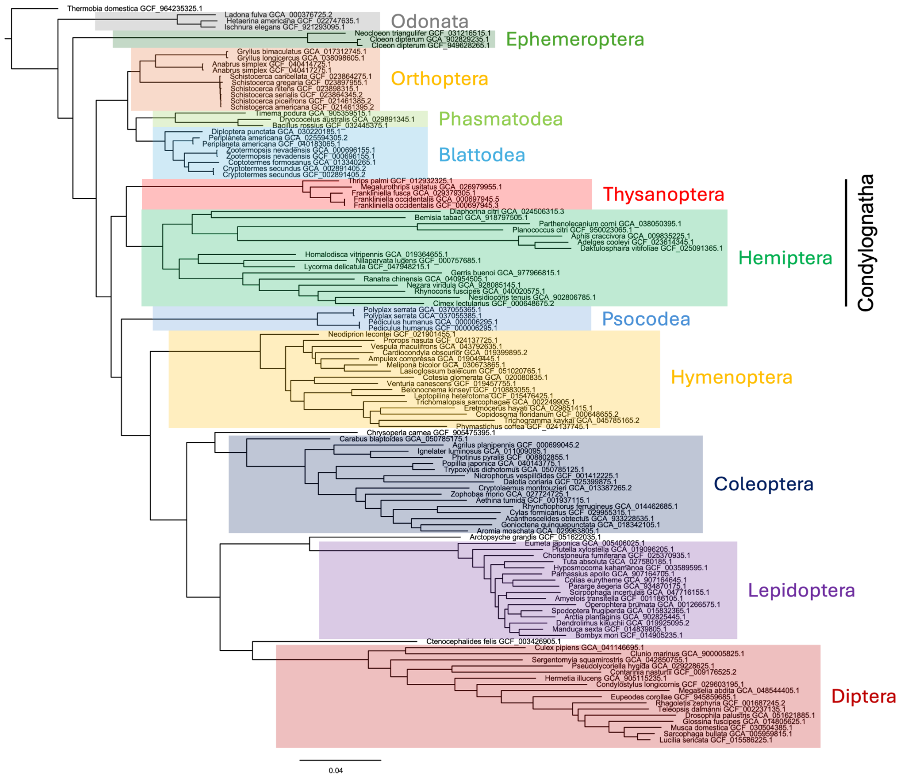

**Figure S7**

Phylogenetic tree inferred with IQ-TREE from the ‘BUSCO taxa’ alignment of class Insecta. The tree was rooted using Thermobia domestica (Zygentoma) as the outgroup, and order names are shown for orders represented by at least two samples.

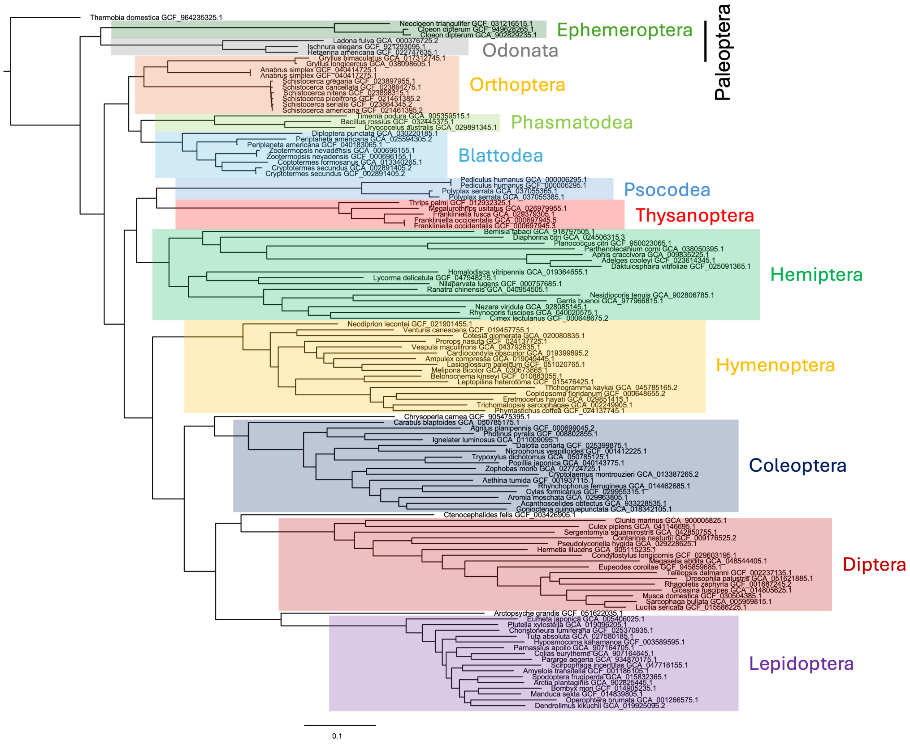

**Figure S8**

IQ-TREE phylogeny of the 190 mammals under the LG+I+G model with 2,000 ultrafast bootstrap replicates. Bootstrap values are shown only for branches with support below 100%.

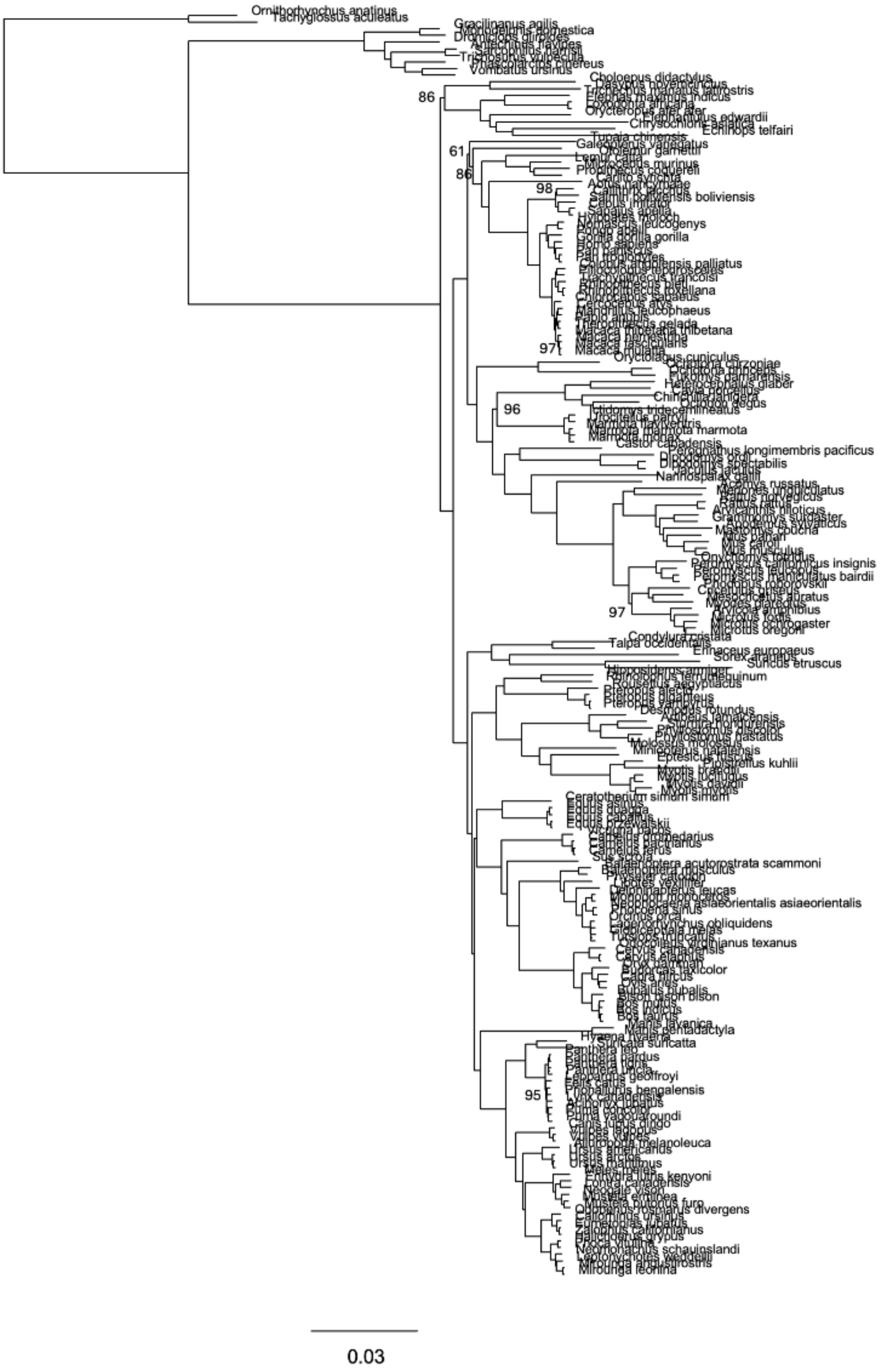

**Figure S9**
IQ-TREE phylogeny of the 190 mammals under the LG+C20+I+G model with 2,000 ultrafast bootstrap replicates. Bootstrap values are shown only for branches with support below 100%.

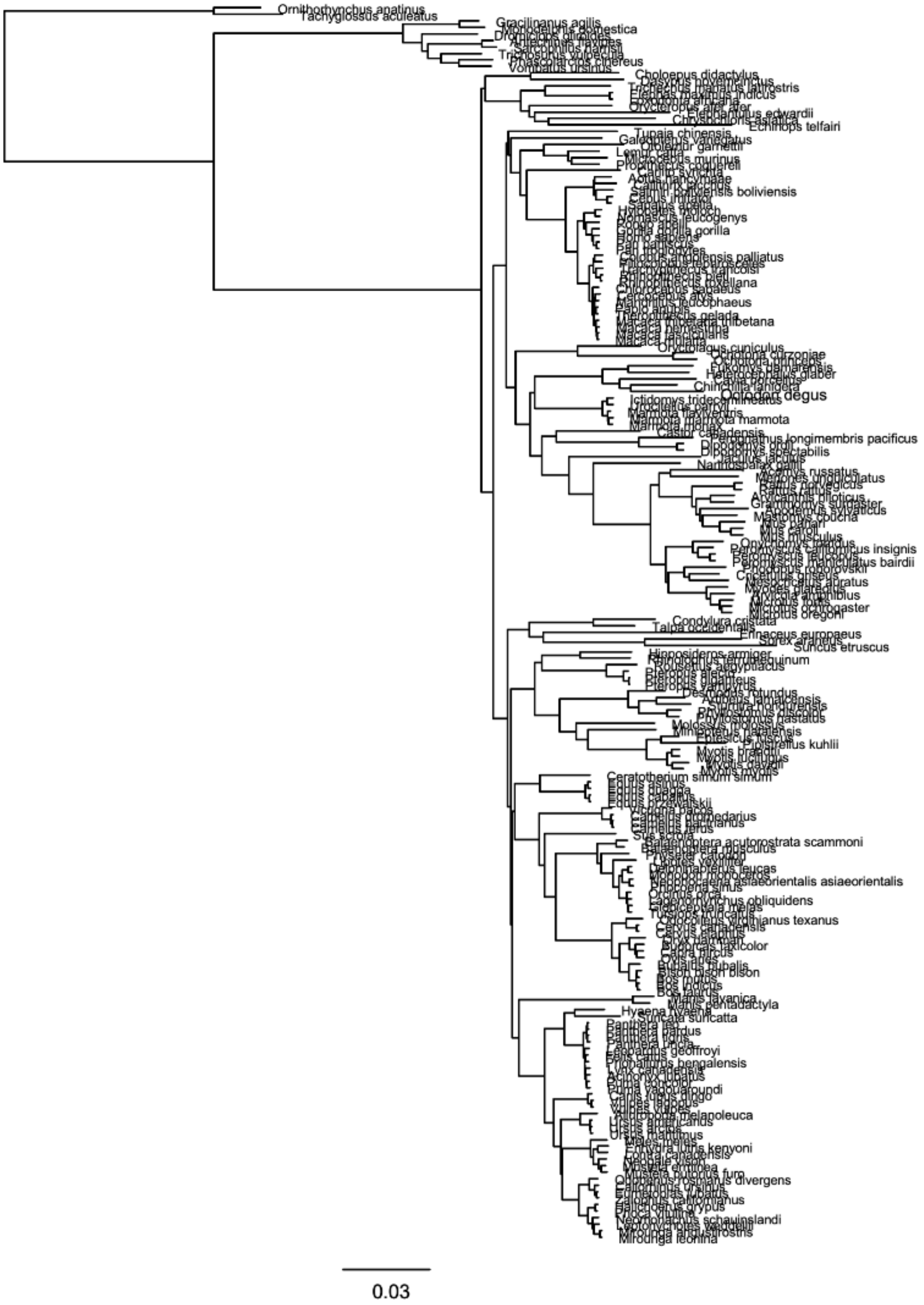

**Figure S10**
Numbers of positions in kamino alignments generated from datasets ranging from 40 to 200 proteomes.

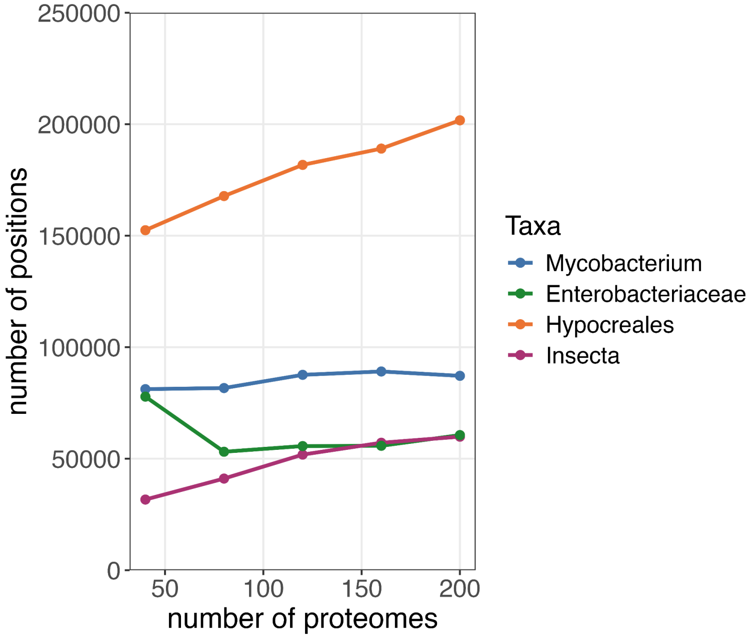


**Figure S11**
Runtime of kamino for the 120-isolate datasets using 1 to 12 CPU threads.

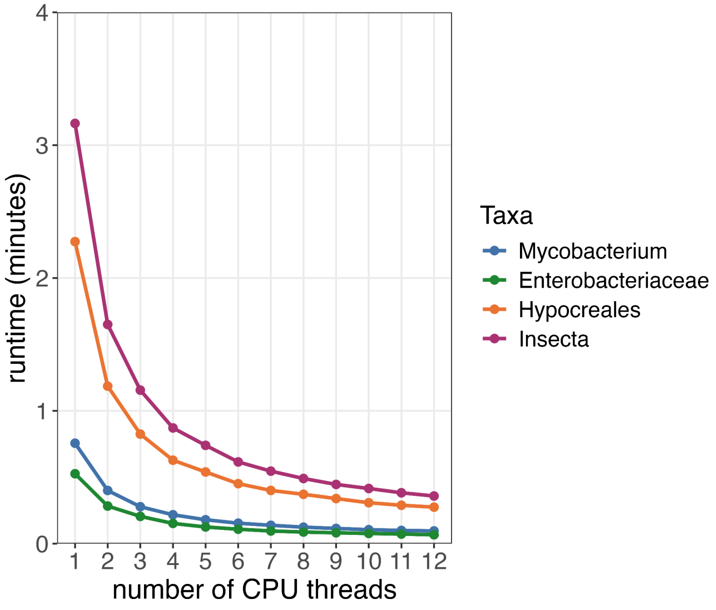
